# Temperature-responsive control of *Pseudomonas aeruginosa* virulence determinants through the stabilization of quorum sensing transcriptional regulator RhlR

**DOI:** 10.1101/2024.05.13.593818

**Authors:** Thays de Oliveira Pereira, Marie-Christine Groleau, Nicolas Doucet, Eric Déziel

## Abstract

The versatile bacterium *Pseudomonas aeruginosa* thrives in diverse environments and is notably recognized for its role as an opportunistic pathogen. In line with its adaptability, *P. aeruginosa* produces various exoproducts crucial for survival and virulence, several of which regulated through quorum sensing (QS). These factors are also regulated in response to environmental cues, such as temperature changes. As a pathogen, *P. aeruginosa* is generally thought to activate its virulence factors at temperatures akin to warm-blooded hosts rather than environmental temperatures. Recent studies elucidated the functional structure of the QS transcriptional regulator RhlR, which depends on the stabilizing effects of its cognate autoinducing ligand, *N*-butanoyl-L-homoserine lactone (C_4_-HSL), and of the moonlighting chaperone PqsE. Given the influence of temperature on biomolecular dynamics, we investigated how it affects RhlR activity using the RhlR-regulated *phzA1* promoter as a proxy. Unexpectedly, we found that RhlR activity is higher at 25°C than at 37°C. This temperature-dependent regulation likely stems from altered RhlR turnover, with the presence of PqsE extending RhlR activity tenfold from its basal level at 37°C to that observed at 25°C. This lower, environmental-like temperature promotes increased affinity between RhlR and C_4_-HSL, a trait significantly compromised in the absence of PqsE. These results suggest that this response depends on the structural integrity of the complex, indicating that temperature functions as an additional regulating and stabilizing factor of RhlR function. Accordingly, lower growth temperature fails to increase the activity of a structurally stabilized version of RhlR. The thermoregulation aspect of RhlR activity and signalling impacts the virulence profile of a mutant unable to produce C_4_-HSL, underscoring its significance in bacterial behaviours and potentially conferring an evolutionary advantage.

**Author Summary:** *Pseudomonas aeruginosa* is recognized for its capacity to colonize vastly different environments, thereby encountering a range of temperatures. The bacterium’s ability to adapt to these settings necessitates finely regulated gene expression. Within this regulatory framework lies quorum sensing (QS), the intercellular communication system used by *P. aeruginosa* to orchestrate the expression of genes responsible for producing diverse exoproducts, including the blue phenazine pyocyanin. RhlR primarily governs the expression of genes required for pyocyanin production, including the *phz1* operon. Unlike other QS regulators, RhlR possesses a distinctive characteristic – in addition to its cognate signalling ligand C_4_-HSL, it depends on the presence of the chaperone-like protein PqsE for stability and activity. This intrinsic instability implies that RhlR may be susceptible to external influences that can modulate its function. Indeed, a lower culture temperature, akin to an environmental-like condition, induces the transcription of the *phz1* operon, used as a proxy for RhlR activity. Using a combination of genetic approaches, we present evidence that this thermoregulation is due to an impact on the stability of the RhlR/C_4_-HSL/PqsE complex. We further show the biological effect of this regulation mechanism in an infection setting, which could underscore a relevant role for other bacterial behaviours.

## Introduction

The Gram-negative bacterium *Pseudomonas aeruginosa* exhibits remarkable adaptability attributed to its extensive genomic repertoire, enabling its survival in various ecological niches. Found in natural habitats, environmental isolates of *P. aeruginosa* are mostly distributed in niches closely related to human activity, including contaminated soils and water reservoirs [1–3]. In addition to its frequent isolation from environmental sites, *P. aeruginosa* is a clinically important opportunistic pathogen, frequently involved in nosocomial infections and posing a particular threat to people with cystic fibrosis (CF), leading to severe morbidity and mortality [4]. Interestingly, *P. aeruginosa* infections extend beyond warm-blooded hosts, encompassing reptiles, insects, and plants [5–8]. Virulence determinants are highly conserved across *P. aeruginosa* isolates regardless of their origin (environmental or clinical), underscoring the importance of these factors in diverse natural environments [9].

The production of virulence determinants in *P. aeruginosa* is primarily regulated at the transcriptional level, ensuring appropriate responses to environmental cues. Several regulatory systems are involved in this regulation, including quorum sensing (QS). QS relies on the production of signalling molecules called autoinducers, which, upon entry into the cells, interact with cognate transcriptional regulators, enabling coordinated responses across the population based on cell density [10, 11]. In *P. aeruginosa*, a network of three interlinked systems traditionally represents the QS circuitry. The *las* system comprises the LasR transcriptional regulator and the cognate acyl-homoserine lactone (AHL) synthase LasI. The latter produces the autoinducer *N*-(3-oxododecanoyl)-L-homoserine lactone (3-oxo-C_12_-HSL), which binds to LasR and activates its transcriptional function [12]. Activation of LasR leads to the transcription of several target genes, including coding for virulence determinants and other QS regulatory elements [13–15]. The *rhl* system is composed of the transcriptional regulator RhlR and the associated synthase RhlI, responsible for synthesizing signal *N*-butanoyl-L-homoserine lactone (C_4_-HSL) [16]. Activated by LasR, the *rhl* system controls the expression of genes related to the production of numerous virulence determinants, including the redox-active blue phenazine pyocyanin [13, 15, 17–20].

In addition to the AHL-based QS systems *las* and *rhl*, the QS regulatory network of *P. aeruginosa* also includes the *pqs* system. Unlike the AHL-based systems, the *pqs* system relies on the production of autoinducers belonging to the 4-hydroxy-2-alkylquinolines family (HAQs). The synthesis of HAQs is activated by the QS transcriptional regulator MvfR (also known as PqsR) [21, 22]. Among the enzymes required for the HAQs synthesis are PqsA-D, encoded by the *pqsABCDE* operon [22–24]. Notably, the PqsE protein, encoded by the last gene in this operon, plays a crucial role in fully activating the *rhl* system. Despite the suggestion of a functional interaction between the *rhl* and *pqs* systems for two decades, it was only recently elucidated [25–31]. PqsE moonlights as a chaperone, enhancing the stability of RhlR and, in conjunction with the cognate signal C_4_-HSL, mediating the complete activation of RhlR targets [28, 29].

For opportunistic pathogens such as *P. aeruginosa*, regulating virulence determinants also involves the capacity to sense and adapt to temperature variations. When this bacterium colonizes the human body, it encounters a temperature environment distinct from aquatic and soil settings. Previous studies have documented the thermoregulation of specific virulence factors in *P. aeruginosa*, including those under the QS control [32–36]. Genes under RhlR control, including those implicated in pyocyanin production, are thermoregulated as the protein level of RhlR varies within the cell according to growth temperatures [37]. The synthesis of phenazines involves enzymes encoded by two paralogous operons, *phzA1-G1* (*phz1*) and *phzA2-G2* (*phz2*) [38]. These enzymes produce phenazine-1-carboxylic acid (PCA), which is further converted into derivates phenazines such as pyocyanin [38]. While some high-throughput gene expression profiling comparing the effects of temperature variations indicate the upregulation of the expression of the *phz* operons at 37°C compared to environmental temperatures [32, 34], some report no such regulation [33, 35].

The dynamics of RhlR activity and the subsequent expression of RhlR-dependent virulence determinants are intricate and extend beyond a mere dependence on RhlR protein concentrations. The distinctive activation of this regulator was uncovered using the transcription of *phz1* as a proxy for RhlR activity [26, 28, 29], and the expression data concurred with crystal structures of the active RhlR/C_4_-HSL/PqsE complex [28].

The intricacies of regulating the *rhl* system likely hold ecological significance for an adaptable bacterium like *P. aeruginosa*. Indeed, the emergence of LasR-defective isolates is a common adaptation of this bacterium in both environmental and clinical contexts [39–41]. In these strains, QS-responsiveness persists due to the activity of RhlR and, consequently, of the *rhl* system as a whole [39, 42–45]. Notably, the production of QS-controlled products, such as pyocyanin, has a positive impact on *P. aeruginosa*’s population dynamics [46–50].

Given that QS controls the expression of several virulence factors, we hypothesized that the production of virulence determinants under RhlR control would be favoured at 37°C, a temperature mimicking human infection conditions. We used phenazine production as a temperature-sensitive determinant to explore the intricate interplay between PqsE, C_4_-HSL, and the expression of RhlR-dependent regulation at selected environmental (25°C) and mammalian body (37°C) temperatures. While our experiments confirmed that RhlR activity is subject to thermoregulation, we unexpectedly observed indications that RhlR activity is higher at an environmental temperature rather than at the mammalian body temperature. Our findings suggest that temperature acts as a third stabilizing factor for RhlR, working synergistically with C_4_-HSL and PqsE to optimally regulate the activity of this transcriptional regulator.

## Material and methods

### Bacterial strains and growth conditions

Bacterial strains and plasmids used in this study are listed in **Tables 1** and **2**, respectively. Bacterial cultures were routinely grown in tryptic soy broth (TSB; BD Difco, Canada) at 37°C in a TC-7 roller drum (New Brunswick, Canada) at 240 rpm or on lysogeny broth (LB; BD Difco, Canada) agar plates. Cultures for gene expression experiments were grown in TSB at 37°C or 25°C in a Multitron Pro incubator (INFORS HT, Switzerland) at 240 rpm. The following concentrations of antibiotics were added when required: for *P. aeruginosa* PA14 tetracycline at 125 µg/ml (solid) or 75 µg/ml (broth), 300 µg/ml carbenicillin, and 100 µg/ml gentamicin. For *Escherichia coli*, 15 µg/ml tetracycline, 100 µg/ml carbenicillin, and 15 µg/ml gentamicin. Irgasan (20 µg/ml) was used as a counterselection agent against *E. coli*.

**Table 1.** Strains used in this study.

| Strain | Lab ID<br># | Relevant genotype or description | Reference |
| --- | --- | --- | --- |
| *P. aeruginosa* |  |  |  |
| PA14 | ED14 | Clinical isolate from a human burn patient UCBPP-PA14 | [51] |
| PA14 $\Delta rhIR$ | ED4406 | PA14 derivate; unmarked in-frame *rhIR* deletion | [52] |
| PA14 $\Delta rhII$ | ED4407 | PA14 derivate; unmarked in-frame *rhII* deletion | This study |
| PA14 $\Delta pqsE$ | ED36 | PA14 derivate; unmarked in-frame *rhII* deletion | [23] |
| PA14 $\Delta rhII \Delta pqsE$ | ED4408 | PA14 derivate; unmarked in-frame double *rhII* and *pqsE* deletion | This study |
| PA14 $\Delta rhII \Delta pqsE \Delta rhIR$ | ED4700 | PA14 derivate; unmarked in-frame triple *rhII*, *pqsE*, and *rhIR* deletion | This study |
| *E. coli* |  |  |  |
| SM10($\lambda pir$) | ED222 | *thi thr leu tonA lacY supE recA::RP4-2-Tc::Mu Km $\lambda pir$* | Lab collection |

**Table 2.**
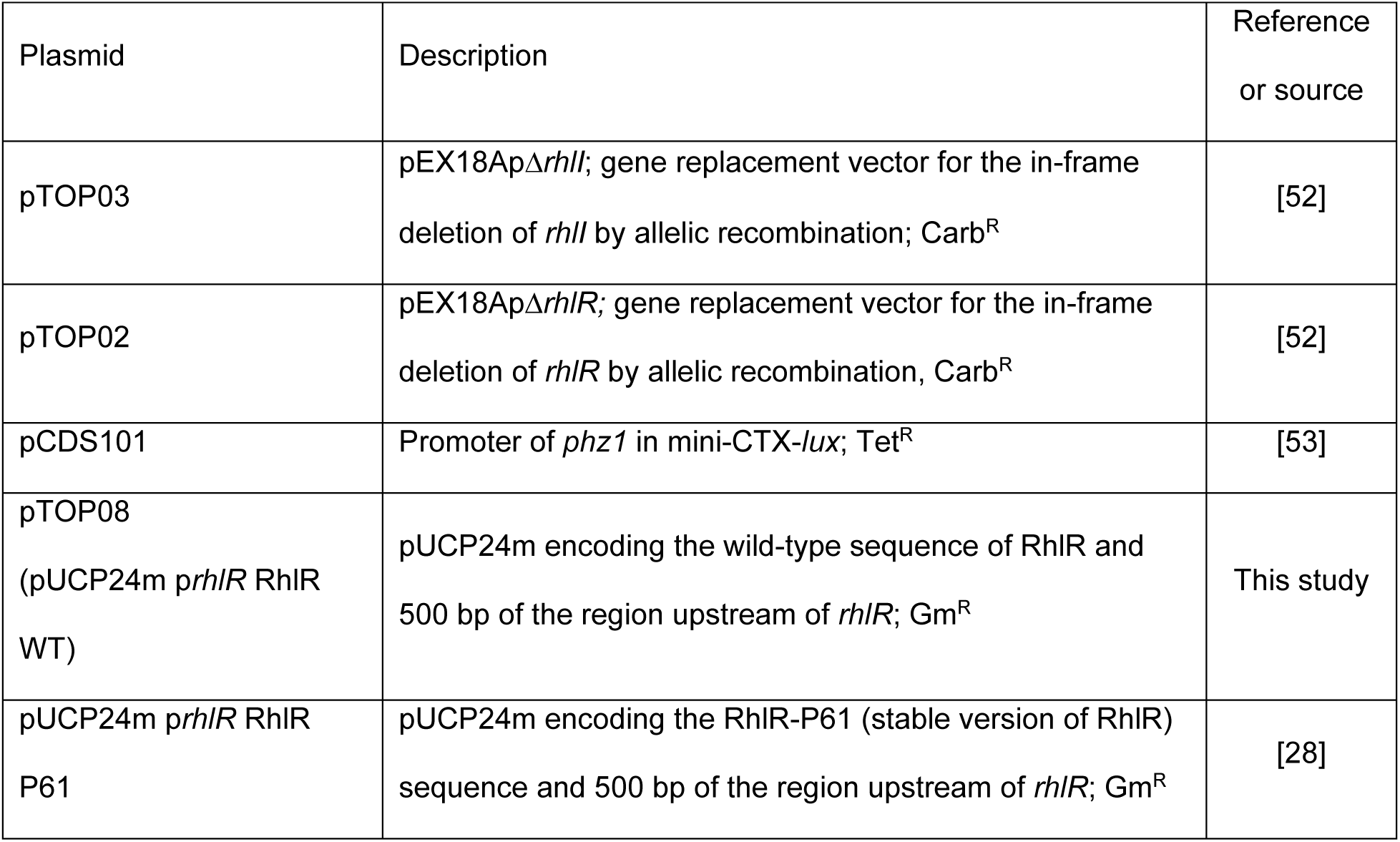
Plasmids used in this study.

| Plasmid | Description | Reference or source |
| --- | --- | --- |
| pTOP03 | pEX18Ap$\Delta rhII$; gene replacement vector for the in-frame deletion of *rhII* by allelic recombination; Carb<sup>R</sup> | [52] |
| pTOP02 | pEX18Ap$\Delta rhIR$; gene replacement vector for the in-frame deletion of *rhIR* by allelic recombination, Carb<sup>R</sup> | [52] |
| pCDS101 | Promoter of *phz1* in mini-CTX-*lux*; Tet<sup>R</sup> | [53] |
| pTOP08<br>(pUCP24m *prhIR* RhIR WT) | pUCP24m encoding the wild-type sequence of RhIR and 500 bp of the region upstream of *rhIR*; Gm<sup>R</sup> | This study |
| pUCP24m *prhIR* RhIR P61 | pUCP24m encoding the RhIR-P61 (stable version of RhIR) sequence and 500 bp of the region upstream of *rhIR*; Gm<sup>R</sup> | [28] |

### Construction of plasmids

pTOP08 (pUCP24m p*rhlR* RhlR WT) was constructed based on pUCP24m p*rhlR* RhlR P61[28]. The latter was linearized by restriction enzymes *Bam*HI and *Hind*III. The p*rhlR* RhlR WT, comprising 500 bp upstream *rhlR* and its coding sequence, was amplified by PCR from PA14 genomic DNA and then cloned into the linearized pUCP24m backbone (**Table S1** for oligonucleotide sequences). pTOP08 was assembled from the purified PCR product and linear vector backbone by employing a seamless strategy of ligation-independent cloning (pEASY® - Uni Seamless Cloning and Assembly Kit, TransGen Biotech Co.).

### Construction of in-frame deletion mutants

Suicide vectors (pTOP02 and pTOP03) were transferred into recipient *P. aeruginosa* strains by conjugation with donor *E. coli* SM10. The recipient merodiploid cells were selected with carbenicillin, and *E. coli* donor cells were counter-selected using Irgasan. Double crossover mutants were isolated by sucrose counter-selection, and PCR confirmed the presence of a mutated allele.

### Construction of chromosomal reporter strains

The *phzA1-lux* reporter from vector pCDS101 was integrated into the *attB* chromosomal site of PA14 and isogenic mutants by conjugation on LB agar plates. Tetracycline was used for selection, and Irgasan was used as a counterselection agent against *E. coli*.

### Luminescence reporter readings

Luminescence was measured using a Cytation 3 multimode plate reader (BioTek Instruments, USA) at several time points during bacterial growth in TSB. The integration time of each reading was one second, a parameter that was used to determine the rate of RhlR activity (*V*_max_). Relative light units (RLU) were normalized by cell density (OD_600_); the latter was measured using a NanoDrop ND100 spectrophotometer (Thermo Fisher Scientific, Canada). When mentioned, C_4_-HSL was added to the indicated concentrations from a stock prepared in high-performance liquid chromatography (HPLC)-grade acetonitrile. Solvent only was added in controls.

### RNA extraction and quantitative reverse transcription-PCR experiments (RT-qPCR)

Total RNA was extracted from PA14 cultures grown in TSB at either 37°C or 25°C using *TransZol* RNA extraction reagent (TransGen Biotech Co.). Cells from three independent cultures were harvested at an OD_600_ of 1.8. After extraction, total RNA underwent two treatments with the TURBO DNA-Free kit (Ambion Life Technologies), following the manufacturer’s guidelines. cDNA was synthesized using iScript reverse transcription supermix (Bio-Rad Laboratories) and used as a template for amplification cycles conducted on a Corbett Life Science Rotor-Gene® 6000 thermal cycler with SsoAdvanced universal SYBR green supermix (Bio-Rad Laboratories). The reference gene was *nadB* [54], and the specific oligonucleotide sequences can be found in **Table S1**. The 2^−ΔΔCT^ formula was applied to determine the relative gene expression [55].

### Galleria mellonella infection

The greater wax moth (*G. mellonella*) larvae were infected by direct injection of *P. aeruginosa* into the last abdominal proleg, with minor adjustments to a previously published protocol [56]. Briefly, bacterial overnight cultures were diluted in fresh TSB to an OD_600_ of 0.05. The cultures were then incubated at 37°C until they reached an OD_600_ of 0.6-0.8. Cultures were then rapidly cooled on ice for 5 min. The bacterial density was then adjusted at a cellular density equivalent to 5×10^3^ CFU mL^-1^ in 10 mM MgSO_4_ solution and maintained on ice until the time of infection. For the infection process, larvae were anesthetized under a gentle carbon dioxide stream and infected by injecting 10 µL of the ice-cold bacterial suspension. Each experimental condition comprised five larvae, and this was replicated with three independent infection groups for each condition, resulting in a total of 15 individuals tested per condition. The data was compared against an infection control group, which received only sterile 10 mM MgSO_4_ solution. Larvae were then incubated at 37°C or 25°C, and morbidity and mortality were monitored for 46 hours to assess the infection score. The entire experiment was conducted twice.

## Results and discussion

### Growth temperature modulates the expression of RhlR-controlled survival determinants

Quorum sensing regulates the transcription of several extracellular products, typically referred to as virulence factors or determinants [57]. These factors are crucial in host infections and bacterial colonization across diverse environments. Therefore, we will adopt the term “survival determinants” to better encompass their broader functions. In *P. aeruginosa*, these survival determinants are predominantly regulated by one of the QS systems, but sometimes by multiple ones, as is the case for *hcnABC* and *lasB* [13, 27, 58]. The production of the redox-active phenazine pyocyanin depends on the expression of the *phz1* and *phz2* operons, which are primarily controlled by the *rhl* QS system [26]; its easily detectable accumulation in cultures has served as a reliable proxy for assessing RhlR activity alongside the expression of the *phz1* operon [26, 28]. The regulatory influence of RhlR on *phz1* transcription is attributed to a *las-rhl* box motif within its promoter region [59]. Importantly, optimal activation by RhlR depends on the involvement of both PqsE and C_4_-HSL [28, 29]. Given these considerations, we used *phz1* transcription as a proxy to assess the activity status of RhlR under two different temperature conditions.

To investigate the influence of temperature on the function of RhlR, we measured the activity of a chromosomal *phzA1-luxCDABE* reporter in the wild-type (WT) *P. aeruginosa* strain PA14. These measurements were carried out at two distinct temperatures: 37°C, mimicking conditions within mammalian hosts, and 25°C, representing an environmental-like temperature. The transcription profile of *phz1* was similar under these conditions, and the expression peak was achieved at an equivalent cell density at both temperatures (i.e., OD_600_ of 1.8) (**Fig. 1**). Remarkably, a threefold increased expression level was measured at 25°C compared to 37°C, indicating that *phz1* transcription is higher at a lower temperature.

**Figure 1.**
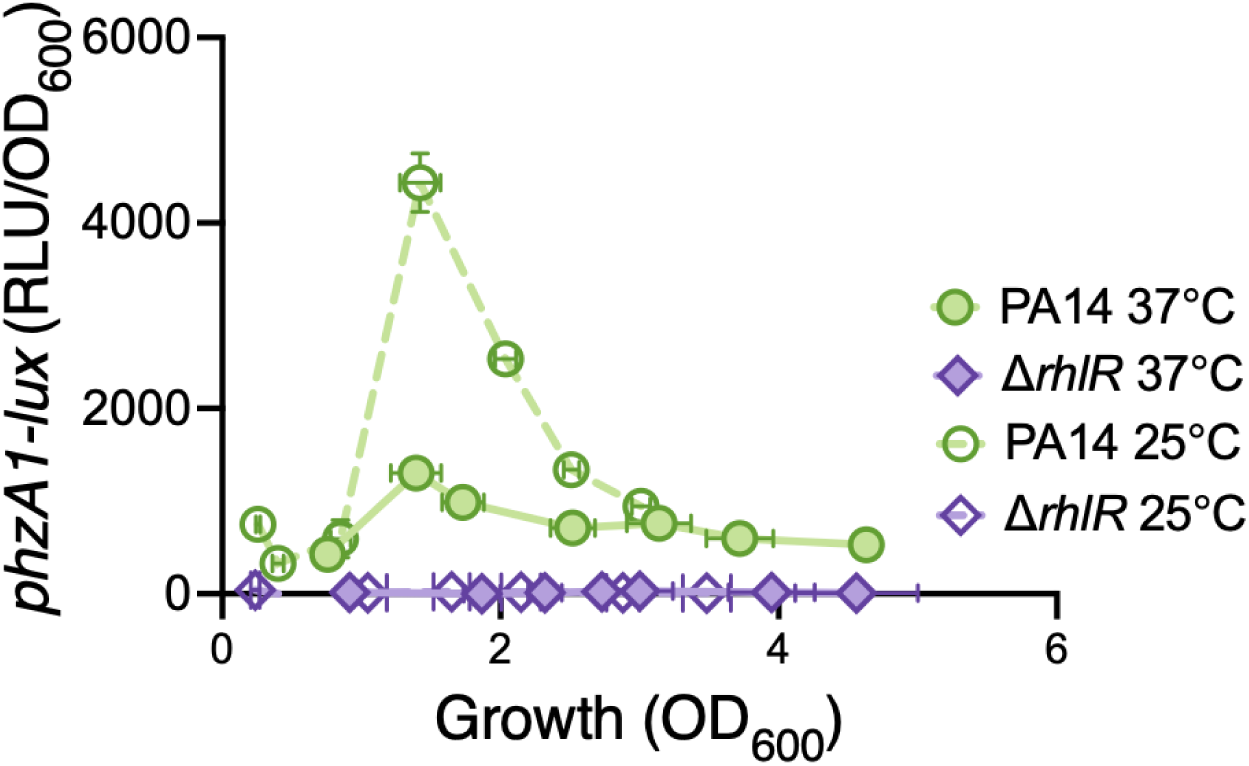
The transcription of *phz1* requires RhlR and is higher at an environmental-like temperature than at body temperature. Luminescence readings (RLU) were used to measure the expression of the chromosomal reporter *phzA1-lux* during growth in TSB of the wild-type PA14 strain and its isogenic *rhlR* mutant (Δ*rhlR*). The cultures were incubated at 37°C (solid symbols and solid lines) and 25°C (open symbols and dotted lines). The values are means ± standard deviation (error bars) from three replicates.

Noteworthy, previous studies have employed the bioluminescence *luxCDABE* reporter to investigate the impact of temperature on gene regulation [60–63]. A 2.5-fold decrease in light production was revealed in bacterial cells grown at 25°C compared to 37°C due to reduced enzymatic activity [60]. Importantly, the relative light unit (RLU) values presented in this study do not account for this 2.5-fold reduction. Consequently, the observed higher *phz1* transcription at 25°C may even be an underestimation. To confirm the role of RhlR in this temperature-responsive behaviour, we conducted the same measurements with an isogenic *rhlR* deletion mutant (Δ*rhlR*). As expected, the transcription of *phz1* was abolished at both temperatures in the absence of RhlR (**Fig. 1**) [26, 64, 65]. This confirmed that the thermoregulation of *phz1* transcription relies on RhlR and suggested that an environmental-like temperature may enhance RhlR expression or its activity.

Based on previous literature that reported an induction of pyocyanin expression at human body temperature [32, 33, 37], we anticipated a similar response. However, our findings diverge from this initial hypothesis. To address this unexpected result, we validated the transcriptional *phz1* reporter by measuring the mRNA levels of *phzA* using RT-qPCR. The measured relative expression of *phzA* was augmented at 25°C compared to 37°C, providing additional evidence that *phz1* transcription is indeed induced at 25°C (**Fig. S1**). This validation underscores the reliability of the *phzA1-lux* reporter to measure the transcription of the *phz1* operon and again established it as a valuable tool for investigating the transcriptional thermoregulation of this RhlR-controlled survival determinant.

One possible reason for the different transcriptional profile of *phzA1* between both temperature conditions is a potential variation in RhlR expression. To explore this possibility, we performed RT-qPCR analyses to assess the transcription levels of *rhlR*. This is necessary because this gene includes four promoters located directly upstream of it and one responsible for driving the expression of a large transcriptional unit containing *rhlA*, *rhlB*, and *rhlR* [66, 67]. No significant differences in *rhlR* mRNA levels were measured between cells grown at 37°C and 25°C (**Fig. 2A**), aligning with previous research comparing *rhlR* transcription from the four proximal promoters under similar conditions [37]. A previously described thermoregulation of RhlR expression relies upon the large transcript, and its presence was characterized under phosphate limitation [37]. Notably, the transcription initiated from *rhlR*’s gene promoters is condition-specific [66]. This alternative transcript has not been described in *P. aeruginosa* cultures grown in rich media, such as TSB. Compared to the previous study, our results suggest that the thermoregulation of *phz1* expression is not linked to altered transcriptional levels of *rhlR* and is thus independent of the large transcriptional unit [37]. Further investigation into alternative mechanisms, including the potential thermoregulation of RhlR activity, is necessary to understand the *phz1* transcriptional profile under different temperatures fully.

**Figure 2.**
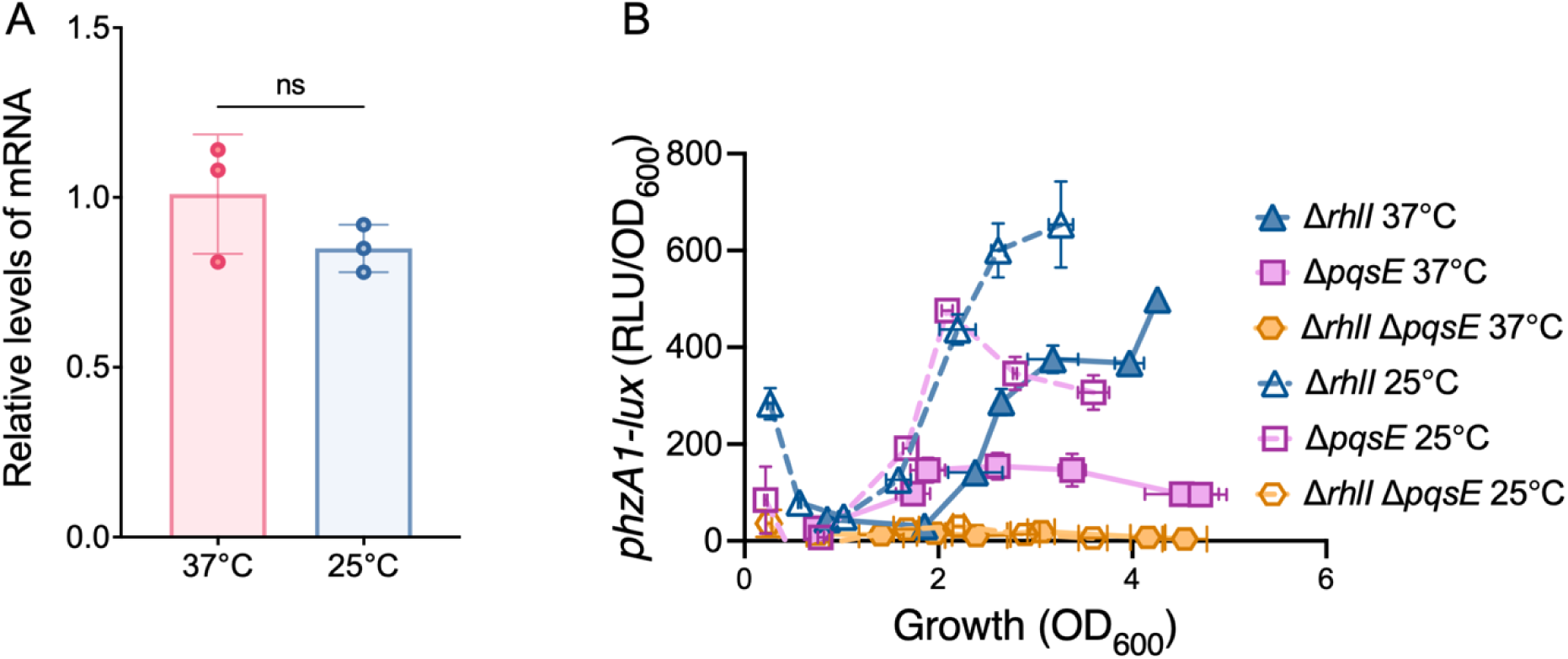
RhlR activity, rather than its expression, mediates the thermoregulation of *phz1* transcription. (A) Expression of *rhlR* was assessed by RT-qPCR from total RNA isolated from wild-type PA14 strain grown in TSB at 37°C (red) and 25°C (blue). The results are presented as relative quantification compared to PA14 grown at 37°C, which was set as 1. The values are means ± standard deviation (error bars) from three biological replicates. An independent t-test was used to quantify statistical significance. ns, non-significant. **(B)** Activity of the chromosomal reporter *phzA1-lux* was measured during growth in TSB in the strains Δ*rhlI*, Δ*pqsE* and the double mutant Δ*rhlI* Δ*pqsE* incubated at 37°C (solid symbols and solid lines) and 25°C (open symbols and dotted lines). The values are means ± standard deviation (error bars) from three replicates.

### C_4_-HSL and PqsE are not both required for RhlR-mediated thermoregulation of gene expression

As full RhlR activity depends on both its autoinducer C_4_-HSL and the presence of PqsE [26–29], we measured *phz1* expression in a *rhlI* mutant, which cannot produce C_4_-HSL, and a *pqsE* mutant, to investigate their impact on thermoregulation of RhlR activity. At 37°C, *phz1* transcription remained significant even when either *rhlI* or *pqsE* were inactive (**Fig. 2B**), consistent with prior studies [26]. Moreover, neither mutant strain produced pyocyanin under these conditions, corroborating previous findings and our results (see **Fig. S2A**) [20, 22, 68]. Interestingly, similar to the wild-type strain, transcription from the *phz1* promoter was higher at 25°C than at 37°C in both mutant backgrounds (**Fig. 2B**). This upregulation even restored pyocyanin production in the Δ*rhlI* mutant but not in the Δ*pqsE* mutant (see **Fig. S2B**), potentially due to the greater dependence of *phz1* transcription on PqsE than C_4_-HSL during the stationary phase. We also note that the absence of *rhlI* led to higher transcription levels at both temperatures, consistent with a greater contribution of PqsE than C_4_-HSL on RhlR function in this system. Despite the differing transcription profiles resulting from the absence of PqsE or C_4_-HSL, these results indicate that the RhlR-driven thermoregulation of *phz1* expression does not necessitate the simultaneous presence of both factors.

To further investigate the role of RhlR’s activity in its thermoregulation, we explored conditions where RhlR function would be lost, presumably in the simultaneous absence of C_4_-HSL and PqsE. Hence, we utilized the Δ*rhlI* Δ*pqsE* mutant background to assess the connection between the thermoregulation of RhlR and its activity. Interestingly, *phz1* transcription was completely abolished in this strain at both temperatures, strengthening the idea that temperature influences RhlR function through these factors.

### Environmental temperature enhances RhlR activity

The requirement for either C_4_-HSL or PqsE in the thermoregulation of RhlR activity on *phz1* transcription indicates that this effect is likely going through the functional structure of RhlR. Indeed, the RhlR homodimer exhibits an inherently unstable fold [28, 69, 70]. This characteristic is common to most LuxR-type proteins [71]. Typically, these proteins adopt stable three-dimensional structures when bound to their cognate autoinducers. However, in contrast to typical LuxR-type proteins, RhlR stability requires the presence of both ligand C_4_-HSL and chaperone-like PqsE [28, 29]. Based on the thermoregulation profile, we hypothesized that lower temperatures resembling environmental conditions would contribute to an increase in the stability of the active RhlR complex compared with 37°C, the temperature of warm-blooded animals. In other words, we postulated that temperature variations could influence the actual stability of the RhlR complex, putatively mitigating the absence of C_4_-HSL and PqsE under certain environmental conditions.

To investigate the predicted dynamic behaviour of active RhlR complex formation under different temperature conditions, we used a strain incapable of producing C_4_-HSL (i.e., Δ*rhlI* mutant). We then introduced a defined concentration of exogenous C_4_-HSL and evaluated *phz1* transcription in cultures grown at 37°C or 25°C. Interestingly, when an equivalent concentration of C_4_-HSL was added to cultures grown at 25°C, *phz1* transcription levels were up to tenfold higher than those observed in cells grown at 37°C (**Fig. 3A**). One possible explanation for this transcriptional behaviour could be that temperature variations modulate the affinity of RhlR towards C_4_-HSL. Increased temperatures could also negatively impact the stability of the active RhlR/C_4_-HSL/PqsE complex, and thus effective concentrations in the cell.

**Figure 3.**
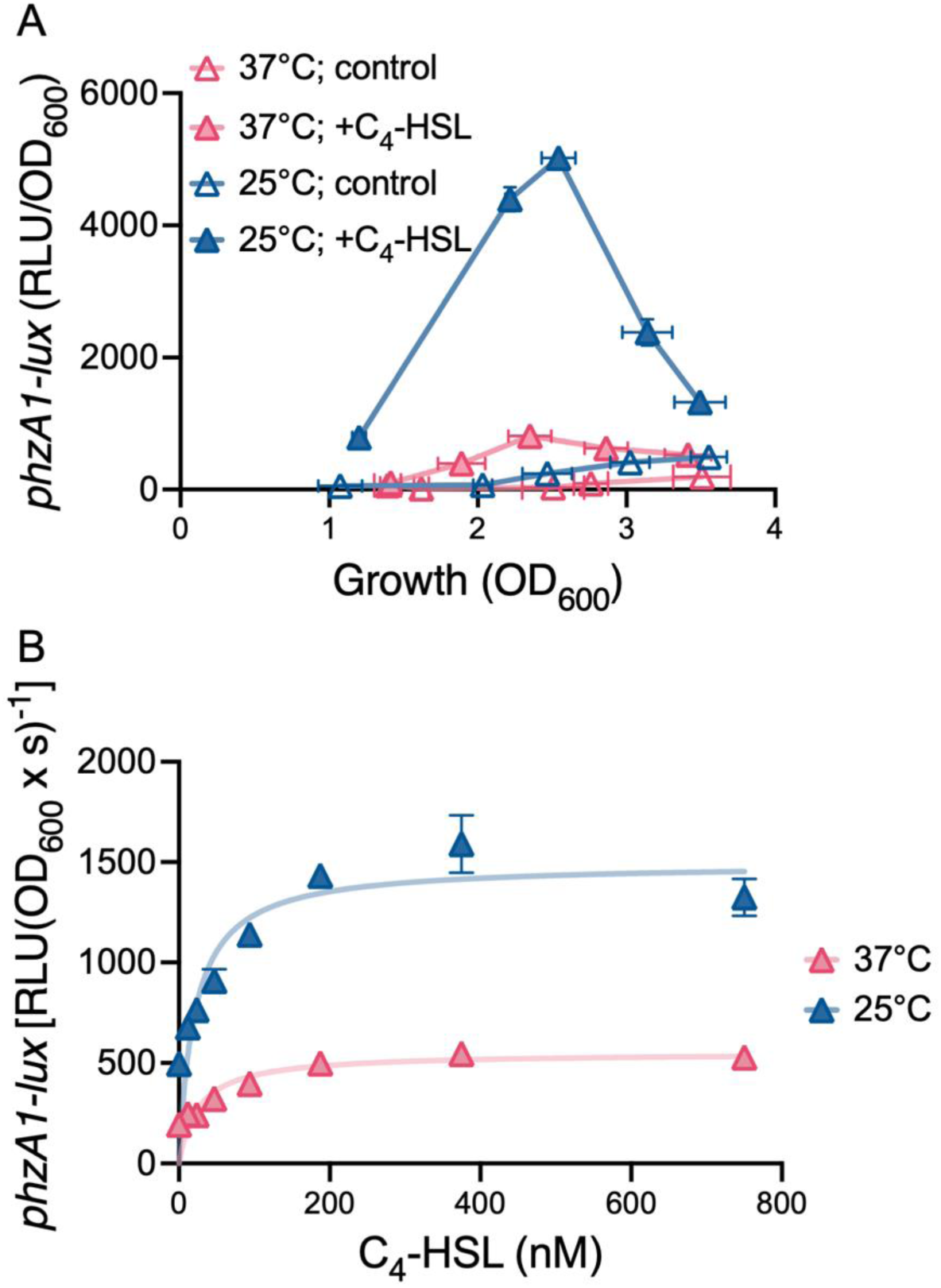
At an environmental temperature (25°C), the maximum activity of RhlR is extended from its basal level at 37°C. (A) Transcription of the chromosomal reporter *phzA1-lux* was measured in the Δ*rhlI* mutant grown in TSB at 37°C (red) and 25°C (blue) in response to supplemented 750 nM C_4_-HSL. Solid symbols represent C_4_-HSL conditions, while open symbols denote control conditions. The values are means ± standard deviation (error bars) from three replicates. **(B)** Increasing concentrations of C_4_-HSL (up to 750 nM) were used to assess their impact on *phzA1-lux* transcription during the stationary growth phase (OD_600_ of 3.4-3.5) at 37°C and 25°C in the Δ*rhlI* background. Data analysis was performed using non-linear regression resembling the Michaelis-Menten enzyme kinetics model (solid lines).

Next, we investigated whether temperature variations could affect the profile of *phz1* transcription. According to our hypothesis, lower concentrations of C_4_-HSL would be required to elicit an equivalent response in cell growth at 25°C relative to 37°C. To investigate this, we assessed *phz1* transcription in response to a series of C_4_-HSL concentrations under defined temperature conditions. The induction profile of *phz1* transcription in response to C_4_-HSL varied during growth phases (**Fig. 3A**). Given this varying response, we compared the transcriptional profiles of cells in equivalent growth phases at 37°C and 25°C. Interestingly, the dose-dependent effect induced by increasing concentrations of C_4_-HSL resulted in a profile akin to steady-state Michaelis-Menten enzyme kinetics, which describes the relationship between substrate concentration and the rate of the corresponding enzymatic reaction [72], predominantly during a stationary growth phase (**Figs. 3B** and **S3**). Although RhlR is not an enzyme and C_4_-HSL is not a catalytic substrate *per se*, the outcome in measured reaction rate (i.e., luminescence as an indicator of transcriptional activity) still follows steady-state kinetics behaviour inherently linked with the enzymatic activity of the corresponding RNA polymerase [73]. This indirect connection between RhlR and RNA polymerase activity could explain the observed pattern, wherein varying concentrations of the C_4_-HSL ligand elicit a corresponding response. Thus, we can draw a parallel between Michaelis-Menten kinetics and our model, whereby the RNA polymerase enzymatic reaction follows RhlR activity in response to C_4_-HSL binding at equilibrium.

From this observation, two relevant kinetic constants were extracted and applied to our model analysis. In a Michaelis-Menten relationship, reaction velocity increases with substrate concentration until it reaches a maximum rate (*V*_max_), indicating that the enzyme is fully saturated by the substrate. Additionally, the affinity of an enzyme for its substrate can be determined by extracting the Michaelis constant (*K*_m_), which represents the substrate concentration at which the enzyme achieves half of its maximum reaction velocity (0.5*V*_max_) [72, 74]. A lower *K*_m_ value indicates higher affinity since the enzyme can achieve half of its maximum velocity at a lower substrate concentration. Conversely, a higher *K*_m_ value suggests lower substrate affinity. Applying this to our biological system, *V*_max_ represents maximal RhlR activity induced by a specific concentration of C_4_-HSL, which in turn allows us to determine the affinity of RhlR for C_4_-HSL (i.e., *K*_m_) at both 37°C and 25°C (**Figs. 3B** and **S3**). Our results show that RhlR affinity for C_4_-HSL remains constant across the temperatures tested, contrary to our hypothesis that lower temperatures would increase the affinity of RhlR for C_4_-HSL (*K*_m_ at 37°C is 20.8 ± 5.8 nM and at 25°C, 25.6 ± 7.2 nM). In contrast, temperature primarily affects the maximum activity of RhlR (i.e., *V*_max_). At 25°C, RhlR exhibits higher activity [*V*_max_ = 1494 ± 89.3 RLU (DO_600_ x s)^-1^] relative to the basal level observed at 37°C [*V*_max_ = 550.5 ± 35.2 RLU (DO_600_ x s)^-1^] (**Fig. 3B**). In simpler terms, regardless of its affinity for C_4_-HSL, the activity of RhlR increases at 25°C and remains unchanged under these temperature conditions. This response is observed throughout bacterial growth phases (**Fig. S3**).

In the context of *in vitro* systems, enzyme concentration is a critical factor influencing *V*_max_ [74]. Thus, the regulation of RhlR concentration under the temperature conditions tested could explain the increased activity of this regulator at 25°C. As mentioned earlier, we did not observe a direct impact of temperature on *rhlR* transcription (**Fig. 2A**). However, it remains unclear whether it affects the post-transcriptional regulation of this gene. Indeed, others have observed a link between these factors, with enhanced RhlR concentration at 37°C compared to a lower temperature [37]. This profile challenges our data, which instead suggests a higher RhlR concentration at 25°C. Nevertheless, it hints at the potential modulation of RhlR concentration by temperature variations. Another factor that could contribute to the increased activity of RhlR at 25°C is an altered turnover number of RhlR under temperature conditions investigated. This change in catalytic rate would result from varying stability of the RhlR/C_4_-HSL/PqsE ternary complex under these conditions. According to this hypothesis, lower temperatures would favour more stable complex formation, allowing it to persist longer at 25°C than at 37°C. Current observations do not provide enough evidence to determine the most probable mechanism for the thermoregulation of RhlR activity.

### In the absence of PqsE, environmental temperature increases the affinity of RhlR for C_4_-HSL

To further understand how temperature influences RhlR activity, we examined *phz1* transcription induced by increasing concentrations of C_4_-HSL in the Δ*rhlI* Δ*pqsE* genetic background. The interaction between the RhlR homodimer and PqsE within the active RhlR complex is critical for stabilizing RhlR [28, 29]. We discussed two alternative explanations for the increased RhlR activity at 25°C and hypothesized that disrupting the ability of RhlR to maintain a stable conformation by the absence of PqsE would help us differentiate between these hypotheses. Notably, akin to the profile obtained with Δ*rhlI*, the exogenous addition of an equivalent concentration of C_4_-HSL resulted in enhanced *phz1* transcription in cultures grown at 25°C compared to those grown at 37°C (**Fig. 4A**). As expected, this response was significantly lower than that elicited by the equivalent concentration of C_4_-HSL in the presence of PqsE, reinforcing the importance of the latter in RhlR activity [26–28]. Due to this partial dependence on PqsE for activation, the concentration of C_4_-HSL required to saturate RhlR activity in the absence of PqsE was considerably higher compared to the *rhlI* mutant at both tested temperatures (for instance, *K*_m_ value shifted from 25.6 ± 7.2 nM to 6.4 ± 0.8 µM C_4_-HSL at 25°C and 20.8 ± 5.8 nM to 33.1 ± 2 µM C_4_-HSL at 37°C) (**Figs. 4B** and **S4**). The relationship between PqsE and the affinity of RhlR for C_4_-HSL has been previously observed in a heterologous system [25], which reinforces our results and underscores the role of PqsE in modulating RhlR activity in response to C_4_-HSL.

**Figure 4.**
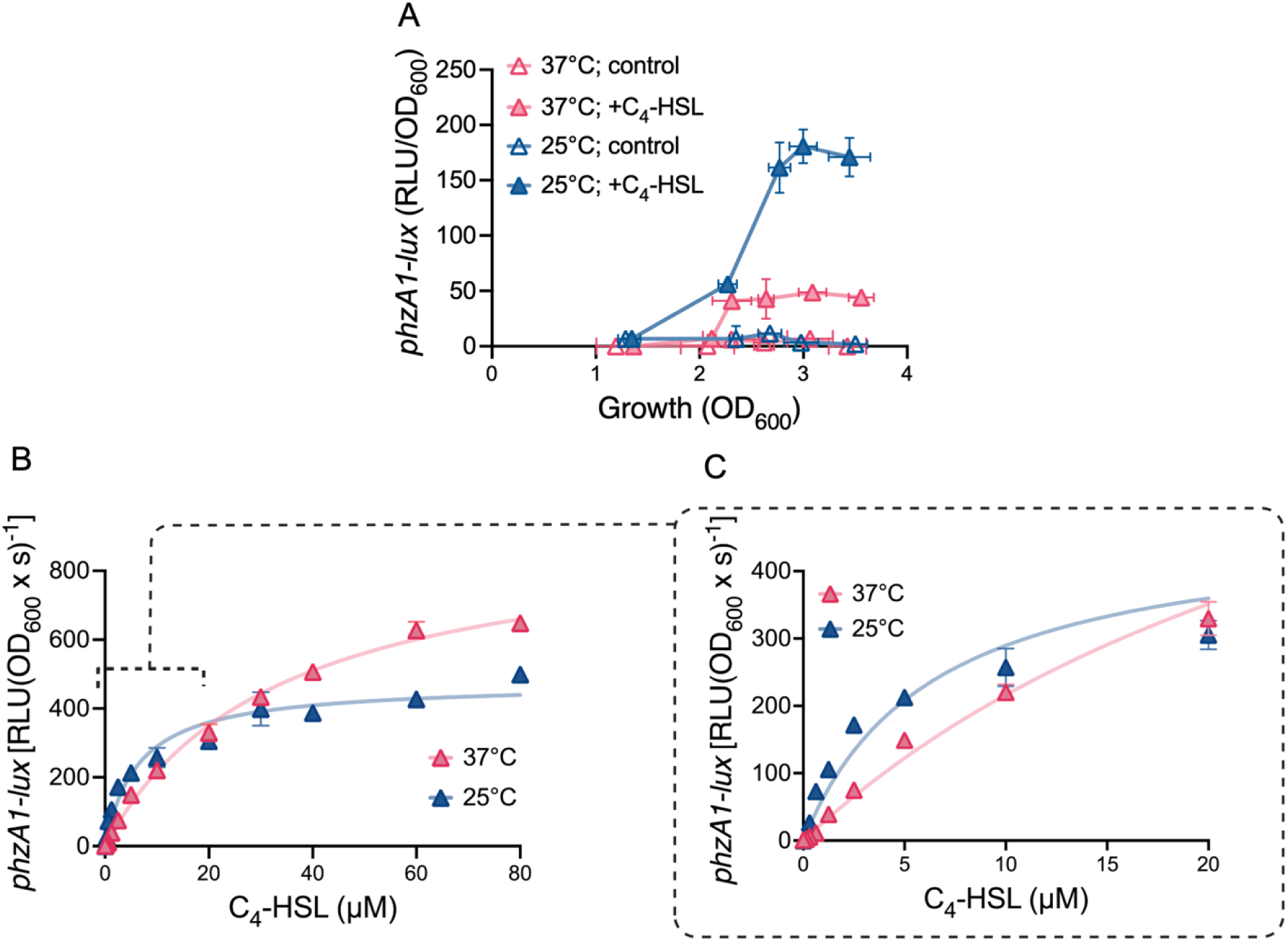
Lower concentrations of C_4_-HSL are required to elicit comparable RhlR activity at environmental temperature in the absence of PqsE. (A) Transcription of the chromosomal reporter *phzA1-lux* was measured in the double mutant Δ*rhlI* Δ*pqsE* grown in TSB at 37°C (red) and 25°C (blue) in response to 750 nM of C_4_-HSL. Solid symbols represent C_4_-HSL conditions, while open symbols denote control conditions. The values are means ± standard deviation (error bars) from three replicates. **(B)** Increasing concentrations of C_4_-HSL (up to 80 µM) were used to assess the impact on *phzA1-lux* transcription during the stationary growth phase (OD_600_ of 3.4-3.5) at 37°C and 25°C in the Δ*rhlI* Δ*pqsE* background. Data analysis was performed using non-linear regression resembling the Michaelis-Menten enzyme kinetics model (solid lines). Panel **(C)** shows an inset of panel B for C_4_-HSL concentrations ranging from 0-20 µM.

By analyzing the saturation profile of *phz1* transcription, we determined the maximum activity of RhlR at both 37°C and 25°C. Interestingly, in the absence of PqsE, we observed that lower temperature did not result in increased activation relative to 37°C, and the maximum activity of RhlR remained relatively stable across growth phases [*V*_max_ = 932.7 ± 25.4 RLU (OD_600_ x s)^-1^ at 37°C and 474.6 ± 15.3 RLU (OD_600_ x s)^-1^ at 25°C]. This thermoregulatory pattern contrasts with that observed in the presence of PqsE (**Figs. 3B** and **S3**), despite employing the same environmental conditions (**Figs. 4B** and **S4**). We reasoned that if the concentration of stable RhlR was the primary driver of the thermoregulated response, as one explanation suggests, we would expect a lower temperature to extend RhlR activity compared to 37°C, similar to the profile of the *rhlI* mutant. However, our data did not support this hypothesis.

Considering these findings, we considered the alternate hypothesis that temperature affects RhlR turnover, possibly through the stability of the complex. The idea is that when the active structure is compromised (e.g., in the absence of PqsE), the stabilizing effect of lower temperature is insufficient to overcome this structural limitation, leading to similar maximal levels of RhlR activity between the temperature conditions tested. Indeed, temperature acts as a stabilizing factor regulating the function of various biological systems. For instance, in prokaryotes such as *Listeria monocytogenes*, a foodborne facultative intracellular pathogen, the formation of a protein complex between MorG and GmaR, triggered at environmental temperatures, modulates the activity of the transcriptional repressor MorG, thereby regulating the transcription of flagellar motility genes [75]. Similar thermoregulatory mechanisms are observed in eukaryotes, as illustrated by *Arabidopsis*. In this plant model, flowering is controlled by a thermoregulated mechanism that stabilizes the RING-finger E3 ligase CONSTITUTIVE PHOTOMORPHOGENIC1 (COP1) protein at lower temperatures [76]. Consistent with these examples, our results suggest that lower temperatures correlate with increased RhlR complex stability in *P. aeruginosa*.

Although temperature does not modulate maximal RhlR activity in the absence of PqsE, it does influence a crucial aspect related to the activity of this regulator. In this genetic context, we observed that RhlR exhibits higher affinity for C_4_-HSL at 25°C (*K*_m_ = 6.4 ± 0.8 µM) than at 37°C (*K*_m_ = 33.1 ± 2 µM) (**Fig. 4B** and **4C**). This indicates that a lower C_4_-HSL concentration is required to achieve maximal RhlR activity at 25°C. Notably, at this temperature, half of the maximum velocity of RhlR is reached with just 6.4 ± 0.8 µM of C_4_-HSL, a concentration within the physiological concentration range of this signalling molecule (ranging from 3.5 µM to 10 µM) [16, 31]. In contrast, the concentration of C_4_-HSL required to elicit an equivalent response increases to 33.1 ± 2 µM at 37°C, which falls outside the physiological concentration range of this molecule in *P. aeruginosa*.

### Mutationally induced RhlR stability disrupts thermoregulation

To validate the model that temperature influences RhlR activity by altering the stability of the active complex, we used a RhlR variant known for its increased intrinsic stability. This variant, RhlR-P61, exhibits 61 amino acid residue substitutions compared to the WT [28]. Importantly, the core regulatory function of RhlR remains unaltered since this variant retains a WT DNA-binding domain [28]. The RhlR-P61 variant activates pyocyanin production independently of C_4_-HSL and PqsE [28]. The involvement of RhlR intrinsic stability in thermally induced responses was investigated by inserting a plasmid encoding either RhlR-P61 or RhlR-WT in a genetic background lacking *rhlI*, *pqsE*, and *rhlR* (triple mutant Δ*rhlI* Δ*pqsE* Δ*rhlR*) and measuring the transcription of the *phzA1-lux* reporter with and without exogenous C_4_-HSL supplementation at both 37°C and 25°C. As expected, in the presence of RhlR-WT, *phz1* transcription remains minimal regardless of the incubation temperature as the strain is deficient in the stabilizing elements C_4_-HSL and PqsE (**Fig. 5A**). The addition of C_4_-HSL induces more robust expression at 25°C, reaffirming the trends observed thus far. In contrast, in the presence of the stabilized RhlR-P61 variant, transcription of *phz1* is restored despite the absence of C_4_-HSL (**Fig. 5B**). This confirms that *phz1* transcription driven by this variant is independent of the presence of C_4_-HSL and PqsE. Notably, the change in incubation temperature did not affect this basal transcription (**Fig. 5B**), indicating that the inherent stability of the RhlR variant remains unaltered by temperature variations. The addition of C_4_-HSL does not result in a significant induction of *phz1* under these conditions, whether at 37°C or 25°C (**Fig. 5**). These findings unequivocally show that the induction of *phz1* expression in the presence of RhlR-WT is responsive to stabilizing factors, such as C_4_-HSL and temperature. The combined presence of both stabilizing factors maximizes the response, and the presence of PqsE further enhances it.

**Figure 5.**
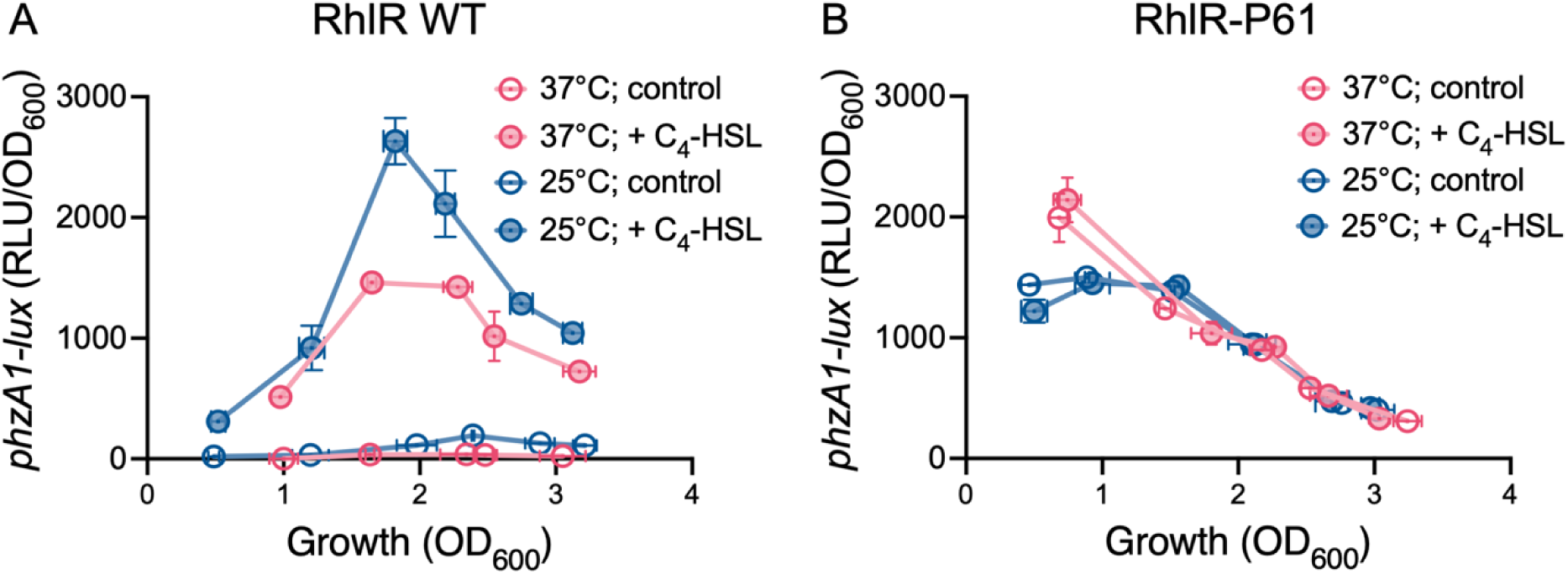
A stabilized variant of RhlR is not sensitive to temperature variations. Transcription of the chromosomal reporter *phzA1-lux* in the triple mutant Δ*rhlI* Δ*pqsE* Δ*rhlR* during growth in TSB. Cultures were incubated at 37°C (red) or 25°C (blue). C_4_-HSL was supplemented at the final concentration of 1.5 μM (solid symbols and lines). Solvent alone (acetonitrile) was used in the controls (open symbols and dotted lines). (A) in the presence of a RhlR-WT encoding plasmid vector. (B) in the presence of a RhlR-P61 encoding plasmid vector.

It is important to note that the sequence cloned into the plasmid used to complement RhlR consists of only 500 nucleotides upstream of the coding region of this regulator. The identification of thermoregulation in the presence of this plasmid reaffirms that this response is independent of the previously described RNA thermometer [37]. This RNA control element, situated upstream of *rhlA*, is not present in the constructed system we employed. In essence, this demonstrates that the thermoregulated responses of the *rhl* system are also influenced by the stability of the active RhlR complex.

### Temperature affects the virulence of a Δ*rhlI* mutant

Considering that *P. aeruginosa* is an opportunistic pathogen and several of its virulence factors are regulated by QS, the implications of the induction of RhlR activity in response to bacterial growth at environmental temperatures are instrumental in understanding its virulence. The functionality of the *rhl* system has been previously associated with bacterial virulence in several infection models [77, 78]. While *rhlR* mutants are unable to produce pyocyanin, a Δ*rhlI* mutant is not. Indeed, this mutant regains the ability to produce this survival determinant at a lower temperature (**Fig. S2**). We hypothesized that a Δ*rhlI* mutant would manifest distinct virulence profiles when exposed to temperatures lower than those resembling mammalian infection conditions. Consequently, we expected that the absence of C_4_-HSL could be compensated by a lower temperature, presumably maintaining the production of RhlR-controlled survival determinants. We expected that the virulence profiles of this mutant would vary between the tested temperatures, while no significant difference in virulence was foreseen for the Δ*rhlR* mutant. To verify this, we needed an infection model capable of operating at different temperatures. Several infection models have been developed for *P. aeruginosa*, each with their unique advantages [5, 7, 79–81]. Among these, the utilization of *Galleria mellonella* larvae, commonly known as the Greater wax moth, provides the necessary flexibility to investigate temperature-dependent variations [56]. According to the expected response, at 37°C, the Δ*rhlR* and Δ*rhlI* mutants exhibit reduced virulence compared to the PA14 wild-type strain (**Fig. 6A**). Fluctuations in temperature significantly influence the virulence profile of *P. aeruginosa*. At 25°C, while the virulence of the Δ*rhlR* mutant towards *G. mellonella* remains significantly different from that of PA14, the virulence of the Δ*rhlI* mutant does not (**Fig. 6B**). This thermo-dependent response aligns with the predicted outcome, indicating that the restoration of RhlR-controlled determinants’ production in the Δ*rhlI* strain at lower temperatures significantly impacts its virulence. The mortality rate of larvae infected with a Δ*rhlR* mutant reinforces the impact of RhlR-controlled survival determinants in this infection model.

**Figure 6.**
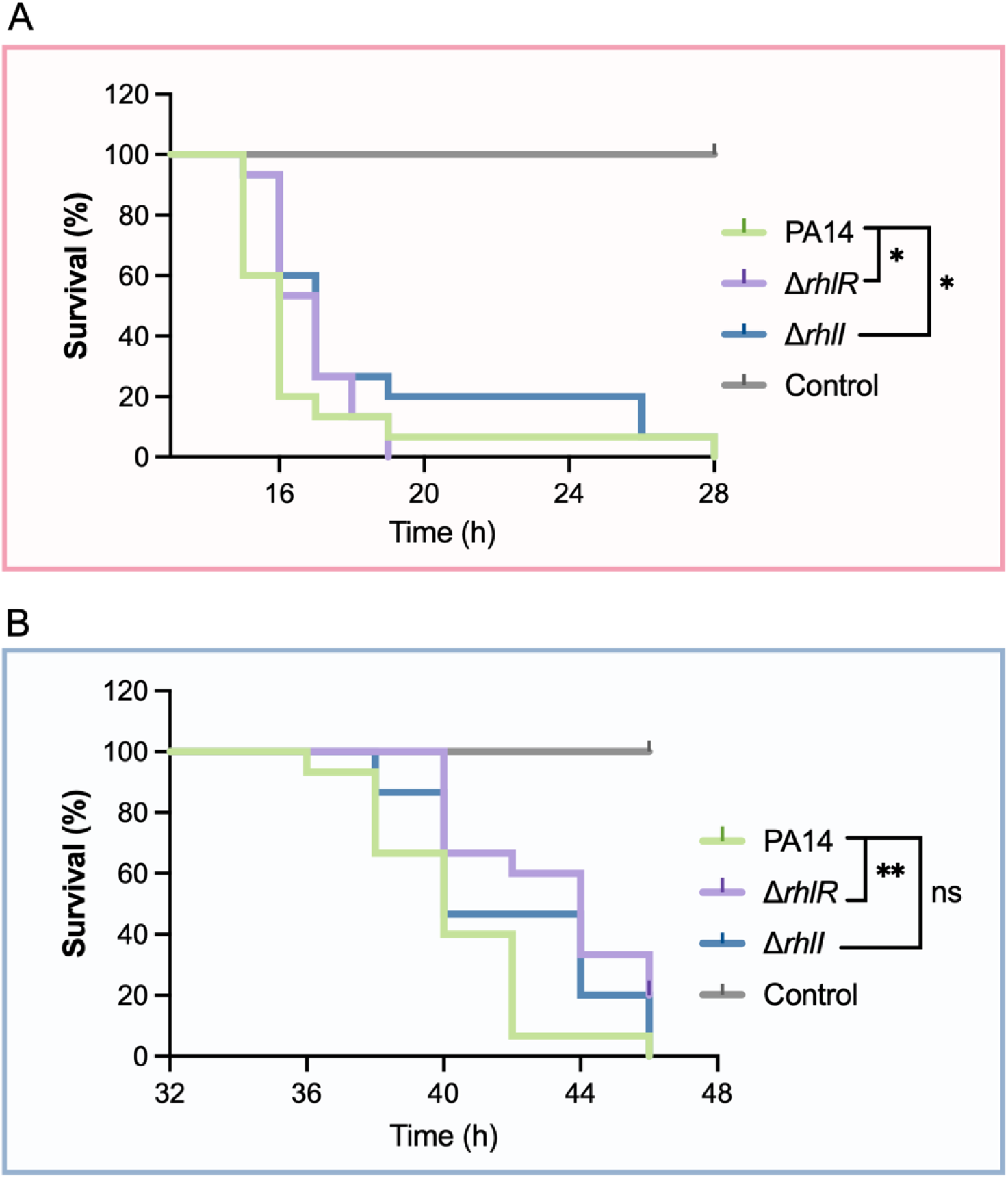
Temperature impacts the virulence profile of a Δ*rhlI* mutant of *P. aeruginosa* toward *G. mellonella*. (A) The great wax larvae were infected with a suspension of 5×10^3^ CFU mL^-1^ (50 bacterial cells per larvae) and incubated at 37°C (top panel, red) or **(B)** at 25°C (bottom panel, blue). Larvae mortality was monitored over time. *N* = 15 larvae per group for each experiment. The experiment was performed independently twice. Statistical significance was determined using Kaplan-Meier analysis and survival curves were compared using Gehan-Breslow-Wilcoxon. ns, nonsignificant, * *P* ≤ 0.05, ** *P* ≤ 0.01.

## Conclusion

*P. aeruginosa* is an adaptable and versatile saprophyte bacterium, isolated from multiple environments, as a free-living bacterium in aquatic and soil habitats and also in association with several hosts. In these environments, similarly to the requirement encountered in human infection settings, the expression of factors under QS control is important.

A paradox within *P. aeruginosa*’s adaptation strategy is the prevalent deficiency in LasR, a critical QS transcriptional regulator. This deficiency is found in approximately 40% of *P. aeruginosa* isolates from various environments and highlights the need for a functioning *rhl* system for QS responsiveness [39, 40]. RhlR, an inherently unstable protein, depends on the presence of both its cognate ligand C_4_-HSL and the chaperone-like protein PqsE for its stability and activity. Within the canonical QS circuitry, LasR is the primary regulator of C_4_-HSL production [26]. Additionally, LasR indirectly stimulates the production of PqsE through MvfR [13, 82]. Consequently, the absence of LasR has a detrimental effect on both key elements required for RhlR function.

We have found that a lower temperature, resembling those encountered in environmental niches, is sufficient to reduce the requirement of RhlR to C_4_-HSL, as it serves as a third stabilizing element of RhlR. This temperature-dependent stabilization exerts a profound influence on the expression of genes under RhlR control, as exemplified by our findings with *phz1*. This thermoregulation could be a strategic adaptation exploited by LasR-deficient cells to maintain QS function, effectively compensating for reduced levels of C_4_-HSL and PqsE. Indeed, this compensatory response extends to situations where C_4_-HSL is absent, as indicated by the continued production of pyocyanin at 25°C.

Our findings hold significant implications, particularly concerning the evolution of *P. aeruginosa* and the *pqsABCDE* operon. Within the *Pseudomonas* genus, *P. aeruginosa* is the most prominent opportunistic pathogen, whereas the other species in the genus are predominantly associated with environmental habitats such as water, soil, or plants [83]. Notably, the *pqs* operon, and thus the associated protein PqsE, is found solely in *P. aeruginosa* within this genus. This observation raises intriguing questions about the role of the *pqs* operon in the transition of *P. aeruginosa* from environmental niches to warm-blooded hosts. The emergence of PqsE may have enabled the activity of the *rhl* system in mammalian infection settings, potentially contributing to the pathogenicity of *P. aeruginosa* in human infections. Interestingly, the sole known homologue of PqsE, HmqE [84, 85], is also present in opportunistic pathogens of the *Burkholderia* genus [86]. This observation hints at a potential conversion of evolutionary traits favouring QS mechanisms in response to environmental cues among opportunistic pathogens. Also interesting, *rhlI* mutations were recently reported to accumulate in naturally occurring LasR-negative isolates [87]. In these cases, the *rhl* system could remain functional at lower temperatures, facilitating bacterial adaptation and enhancing virulence, particularly in non-warm-blooded hosts. These findings underscore our work’s ecological and clinical relevance, emphasizing the remarkable adaptability of *P. aeruginosa*.

## Acknowledgments

We thank Wulf Blankenfeldt from the Helmholtz Centre for Infection Research and the Technical University of Braunschweig for generously providing materials that greatly contributed to the progress of this research.

## Supporting information captions

Figure S1. **Thermoregulation of *phzA* transcription.** Gene expression was assessed through RT-qPCR, utilizing total RNA isolated from wild-type PA14 strain cultures grown in TSB at 37°C (red) and 25°C (blue). Cultures were harvested at an OD_600_ of 1.8 based on the transcriptional profile of *phzA1* (see Figure 1). The results are presented as relative quantification compared to PA14 grown at 37°C, which was set as 1. *phzA1* and *phzA2* have nearly identical sequences and cannot be differentiated. The values are means ± standard deviation (error bars) from three biological replicates. An independent t-test was used to quantify statistical significance. ** *P* ≤ 0.01.

**Figure S2. Thermoregulation of pyocyanin production.** Pyocyanin was chloroform extracted and quantified in the PA14 strain and those with impaired *rhl* system: Δ*rhlR*, Δ*rhlI*, Δ*pqsE*, and the double mutant Δ*rhlI* Δ*pqsE*. These strains were cultivated in TSB and incubated at either 37°C (red, left panel) or 25°C (blue, right panel) for 24h. The values are means ± standard deviation (error bars) from three biological replicates. An independent t-test was used to quantify statistical significance. **** *P* ≤ 0.0001; ns, non-significant.

**Figure S3. The transcriptional profile of *phz1* in response to C_4_-HSL varies with population density and is consistently higher at 25°C compared to 37°C.** Increasing concentrations of C_4_-HSL (up to 750 nM) were used to assess their impact on *phzA1-lux* transcription during bacterial growth in TSB in the Δ*rhlI* mutant. Response profiles were compared at equivalent cell densities, corresponding to growth at 37°C (red) and 25°C (blue). Data analysis was performed using non-linear regression resembling the Michaelis-Menten enzyme kinetics model (solid lines), which requires the saturation of the response and could not be estimated for (A). The response profile varied with the growth phase, with maximum RhlR activity (*V*_max_) tending to be higher at 25°C than at 37°C. (A) Optical density (OD_600_) of 1.2, with not determined *V*_max_ and *K*_m_ for both tested temperatures. (B) OD_600_ of 2.5 and *V*_max_= 1283 ± 65 RLU (DO_600_ x s)^-1^ and *K*_m_= 427.6 ± 43 nM at 37°C, and *V*_max_= 6554 ± 225 RLU (DO_600_ x s)^-1^ and *K*_m_= 246 ± 20 nM at 25°C. (C) OD_600_ of 2.9 and *V*_max_= 699 ± 30.5 RLU (DO_600_ x s)^-1^ and *K*_m_= 90.4 ± 12.5 nM at 37°C, and *V*_max_= 2437 ± 108.2 RLU (DO_600_ x s)^-1^ and *K*_m_= 48.4 ± 8 nM at 25°C. The accurate determination of *V*_max_ (maximum RhlR activity) and *K*_m_ (RhlR affinity for C_4_-HSL) requires a saturated profile. The response profile tends to become increasingly saturated throughout growth, with full saturation observed only during the stationary phase, as depicted in Figure 3B at an OD_600_ of 3.4-3.5. Thus, the curves presented here solely exhibit a behavioural tendency of RhlR activity and its affinity in response to the tested temperatures.

**Figure S4. The affinity of RhlR for C_4_-HSL is temperature-dependent in the absence of PqsE.** Increasing concentrations of C_4_-HSL (up to 80 µM) were used to assess their impact on *phzA1-lux* transcription during bacterial growth in TSB in the Δ*rhlI* Δ*pqsE* genetic background. Response profiles were compared at equivalent cell densities, corresponding to growth at 37°C (red) and 25°C (blue). Data analysis was performed using non-linear regression resembling the Michaelis-Menten enzyme kinetics model (solid lines). **(A)** Optical density (OD_600_) of 1.4 and *V*_max_= 138.7 ± 61 RLU (DO_600_ x s)^-1^ and *K*_m_= 150.8 ± 90.9 µM at 37°C, and *V*_max_= 313.5 ± 40.4 RLU (DO_600_ x s)^-1^ and *K*_m_= 80 ± 16.9 µM at 25°C. **(B)** OD_600_ of 2.3 and *V*_max_= 641.5 ± 53.3 RLU (DO_600_ x s)^-1^ and *K*_m_= 48.4 ± 7.9 µM at 37°C, and *V*_max_= 2093 ± 236.2 RLU (DO_600_ x s)^-1^ and *K*_m_= 106.5 ± 18 µM at 25°C. **(C)** OD_600_ of 2.8 and *V*_max_= 888.1 ± 78.9 RLU (DO_600_ x s)^-1^ and *K*_m_= 64.9 ± 10.1 µM at 37°C, and *V*_max_= 816.4 ± 34.8 RLU (DO_600_ x s)^-1^ and *K*_m_= 14.5 ± 1.9 µM at 25°C. The accurate determination of *V*_max_ (maximum RhlR activity) and *K*_m_ (RhlR affinity for C_4_-HSL) requires a saturated profile. The response profile tends to become increasingly saturated throughout growth, with full saturation observed only during the stationary phase, as depicted in Figure 4B at an OD_600_ of 3.4-3.5. Thus, the curves presented here solely exhibit a behavioural tendency of RhlR activity and its affinity in response to the tested temperatures.

